# Information theoretic measures of neural and behavioural coupling predict representational drift

**DOI:** 10.1101/2025.05.12.653436

**Authors:** Kristine Heiney, Mónika Józsa, Michael E. Rule, Henning Sprekeler, Stefano Nichele, Timothy O’Leary

**Affiliations:** Department of Engineering, University of Cambridge, Cambridge, United Kingdom; Department of Computer Science, Norwegian University of Science and Technology, Trondheim, Norway; Department of Computer Science, Oslo Metropolitan University, Oslo, Norway; Modelling of Cognitive Processes, Technical University of Berlin, Berlin, Germany; School of Engineering Mathematics and Technology, University of Bristol, Bristol, United Kingdom; Department of Computer Science and Communication, Østfold University College, Halden, Norway

## Abstract

In many parts of the brain, population tuning to stimuli and behaviour gradually changes over the course of days to weeks in a phenomenon known as representational drift. The tuning stability of individual cells varies over the population, and it remains unclear what drives this heterogeneity. We investigate how a neuron’s tuning stability relates to its shared variability with other neurons in the population using two published datasets from posterior parietal cortex and visual cortex. We quantified the contribution of pairwise interactions to behaviour or stimulus encoding by partial information decomposition, which breaks down the mutual information between the pairwise neural activity and the external variable into components uniquely provided by each neuron and by their interactions. Information shared by the two neurons is termed ‘redundant’, and information requiring knowledge of the state of both neurons is termed ‘synergistic’. We found that a neuron’s tuning stability is positively correlated with the strength of its average pairwise redundancy with the population, and that these high-redundancy neurons also tend to show high average pairwise synergy. We hypothesize that subpopulations of neurons show greater stability because they are tuned to salient features common across multiple tasks. Regardless of the mechanistic implications of our work, the stability–redundancy relationship may support improved longitudinal neural decoding in technology that has to track population dynamics over time, such as brain–machine interfaces.

**Author summary:** Activity in the brain represents information about the outside world and how we interact with it. Recent evidence shows that these representations slowly change day to day, while memories and learned behaviours stay stable. Individual neurons change their relationship to external variables at different rates, and we explore how interactions with other neurons in the population relates to this neuron-to-neuron variability. We find that more stable neurons tend to share information about external variables with many other neurons. Our results suggest there are constraints on how representations can change over time, and that these constraints are exhibited in shared fluctuations in activity among neurons in the population.

## Introduction

Recent experimental observations of activity in brain regions crucial for processing certain stimuli or behaviours find that populations of neurons change their tuning gradually over days. In a number of cases this occurs in the absence of overt task learning [1–5]. The rate of this ‘representational drift’ varies from neuron to neuron, with some maintaining stable tuning over days and others showing more volatile responses [6–8].

It is natural to ask what determines the magnitude of drift, and whether features of a neuron’s relationship to the task and to other neurons in the population predict the volatility of tuning. Variability in drift rate is related to activity level [7], tuning precision [9], and importance in behavioural decoding [8], but these factors do not entirely explain the population heterogeneity. More recent experimental work has shown that the rate of drift depends on the features of the represented stimulus [4], stimulus familiarity, and the frequency of exposure [2]. Variability in behaviour also contributes to the amount of measured drift in stimulus representation [10]. These findings suggest that a neuron’s representations of other variables in the task space contribute to the stability of its tuning to the target variable.

Other work has suggested that a neuron’s interactions with the population may drive plasticity in neural representations [11]. Sweeney and Clopath [12] have shown that in a network model consisting of neurons with diverse learning rates, faster learners show greater population coupling. In contrast, Sheintuch et al. [9] have shown that in hippocampus, the formation of functionally connected assemblies in CA3 is associated with less drift. The link between population interactions and representational drift thus remains an open question. Furthermore, the reconfiguration of the population code impacts many approaches to studying neural correlates of behaviour, including technologies such as brain–machine interfaces [13]. Being able to predict which couplings—among and between neurons and behaviours—are likely to remain stable would enable an improved capacity to track the neural code as the population drifts.

In this study, we consider how interactions among neurons in a population influence their rates of drift. Why do some neurons in the population exhibit greater tuning stability than others? Specifically, we asked whether the amount of redundancy, in an information-theoretic sense, is predictive of a neuron’s tendency to drift. Intuitively, redundancy measures how much a neuron’s firing statistics explain behavioural variability, relative to contribution of other neurons in the population. This relative measure is accompanied by synergy, which quantifies the extent to which pairs of neurons encode complementary aspects of behaviour. Together, these measures account for how behavioural encoding is shared among neurons in a population.

Redundancy may provide insight into constraints on how population representations can change. Spare degrees of freedom in a mapping between neural activity and behaviour should permit changes in a neuron’s tuning curve without entailing a change in behaviour. We hypothesise that neurons with high redundant coupling to the population are tuned to a latent task or stimulus feature that is salient across many contexts and thus their activity is more tightly constrained than for other neurons. These neurons may maintain their tuning via the shared connections underlying their redundant activity providing channels for corrective feedback.

### Information decomposition as a measure of communication

The phenomenon of representational drift centers on the tuning curve: each neuron’s activity is considered in how it relates, on average, to the behaviour or stimulus of interest, and it is said to drift when this relationship changes. However, this perspective neglects communication among neurons in its consideration of only the relationship between the external variable and each single neuron. Furthermore, the tuning curve explicitly neglects trial-to-trial variability by averaging it away. A view of population interactions on a trial-by-trial basis can give more insight into how information is distributed through the population and where each neuron lies in this distributed code.

A common feature of representational drift is that neurons change their tuning in a variety of ways. Our focus in this paper is on possible sources of heterogeneous tuning stability from the perspective of information encoding. We approach this question by reanalysing calcium fluorescence data from Driscoll et al. [1] and Marks et al. [4].

Schematics of the two experimental setups are shown in Fig 1(a). Driscoll et al. [1] recorded from posterior parietal cortex (PPC) daily over weeks as mice navigated a virtual T-maze based on a visual cue given at the start of the maze. Marks et al. [4] recorded primary visual cortex (V1) as mice viewed two types of visual stimulus, passive drifting gratings (PDG) and a movie clip (MOV), weekly over 5–7 weeks. In both experiments, neurons in the population showed heterogeneous stability in their tuning to external variables (Fig 1(b)). Here we examine how this neuron-to-neuron variability in representational drift relates to information shared among neurons.

**Fig 1.**
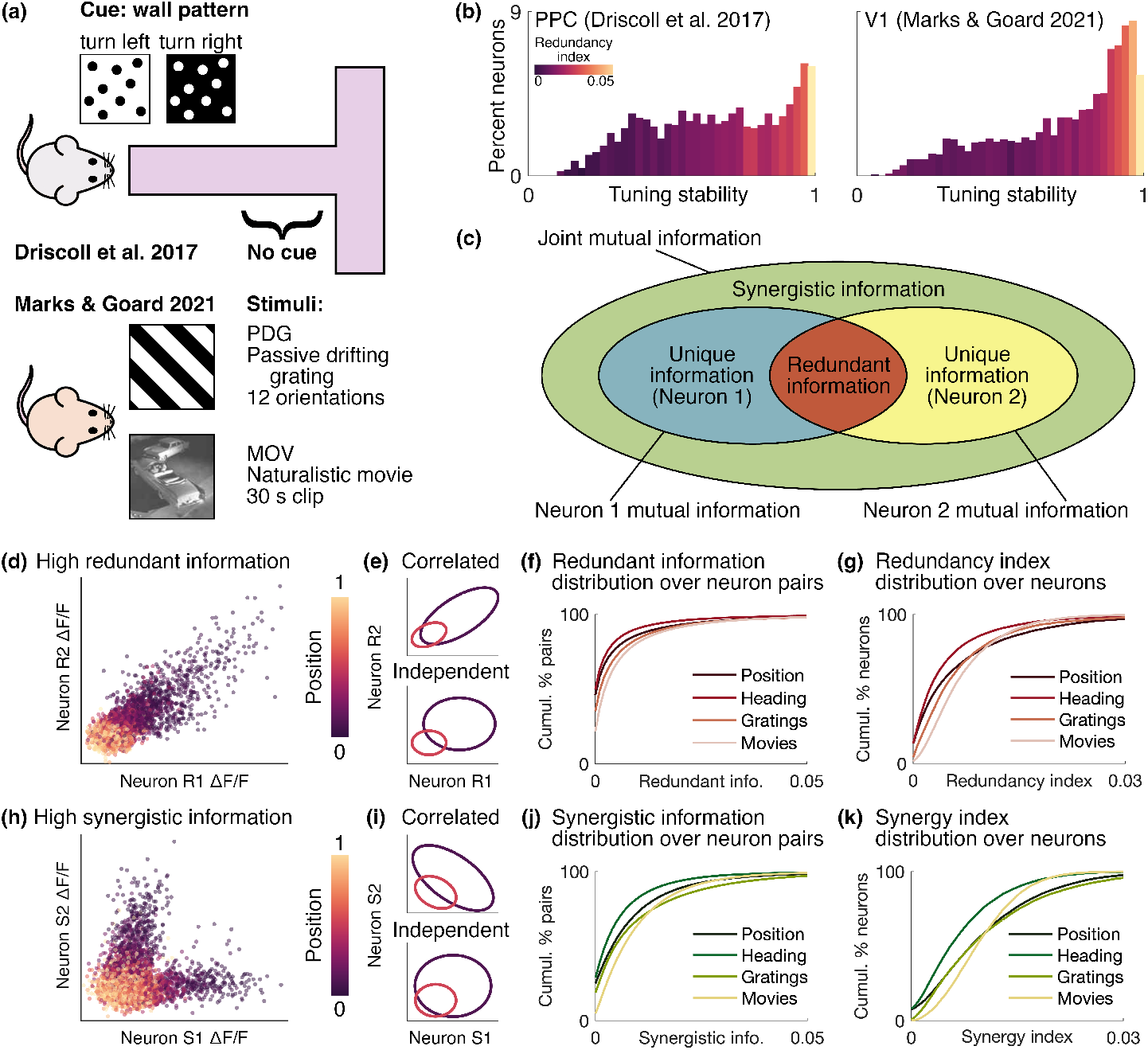
Neural correlations contribute to information encoding of behaviour or stimulus. **(a)** We reanalysed data from mouse posterior parietal cortex (PPC) during maze navigation [1] and mouse primary visual cortex (V1) during presentation of artificial (PDG) and naturalistic (MOV) stimuli [4]. **(b)** Histograms of tuning stability (Eq 4) in the two datasets. Each stability bin is coloured according to the redundancy index averaged over all cells in the bin (see Eq 3). Note that the stability is computed on different time scales between the two datasets. **(c)** Schematic of partial information decomposition. **(d)** Example neuron pair with high redundancy from one mouse on one session [1]. ΔF/F data points are coloured according to the position in the maze. **(e)** Contours of Gaussian distributions fit to the data in **(d)**. We binned the position is binned into 7 bins and fit covariance matrices to bins 1 and 4. The contours in the top plot show these fits, and those in the bottom plot show covariance matrices with the covariance set to 0, corresponding to the neurons acting independent of each other. **(f)** Cumulative distribution of the redundant information over all neuron pairs for each behavior or stimulus. **(g)** Same as **(f)** for redundancy index over all neurons. **(h)–(k)** Same as **(d)–(g)** for synergy.

In this study, we considered neural representations from an information theoretical perspective. This section gives an overview of what this perspective offers; details of the mathematical formulation and the methods used to apply these calculations to empirical data can be found in the methods. In an information theoretical context, a neuron’s tuning can be characterised by the mutual information *I* it shares with the external variable. Mutual information quantifies the extent to which knowledge of one variable reduces uncertainty about the state of another variable. The mutual information between a neuron’s activity *x* and an external variable *u* is given by

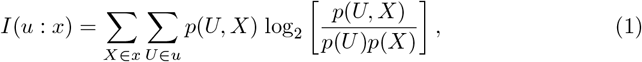

where *x* and *u* are discretised into state bins *X* and *U* and *p*(·) is the probability of a state or joint state occurring. Typically mutual information is considered on a single-neuron basis, but it can be expanded to encompass the information shared between the external variable and a set of multiple neurons. In practice, this presents a statistical challenge, as the number of states that can be taken on by a population increases exponentially with the number of neurons. However, computing the mutual information between the joint activity of a pair of neurons and the external variable is computationally feasible, and it provides insights into how each pair interacts to encode the external variable.

Partial information decomposition (PID) breaks down the mutual information between pairwise neural activity and the external variable into components arising from each neuron independently and from their interactions (Fig 1(c)). In this view, the joint mutual information for a pair of neurons {*x*_1_, *x*_2_} is decomposed into four terms, as

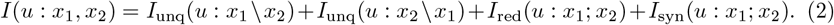

The first two terms here are the information *I*_unq_ provided uniquely by each neuron independent of the other {*x*_*i*_ \ *x*_*j*_}, and the latter two relate to the interactions between the neurons {*x*_*i*_; *x*_*j*_}. The third term is the redundant information *I*_red_, which is the information shared by the two neurons; in an extreme example where the two neurons were copies of one another, the joint information would be entirely composed of redundant information. The fourth term is the synergistic or complementary information *I*_syn_, which is provided only by knowing the state of both neurons together; an example of a synergistic relationship would be an XOR gate, *u* = *x*_1_ ⊕ *x*_2_, with *u, x*_1_, and *x*_2_ binary, where the state of *u* is only known if both *x*_1_ and *x*_2_ are known.

A number of circuit arrangements can give rise to the statistical signatures of redundancy and synergy. For example, a pair of neurons with shared tuning to a second external variable *v* would show high redundancy, as would a pair of neurons that mutually excite one another. In contrast, a pair of neurons with dissimilar tuning to *v* and a mutually inhibiting pair would show high synergy.

We applied PID to the two datasets and calculated the decomposition terms in Eq 2 for all pairs of neurons in a population during each experimental session. We computed estimates of these terms using the definitions given by Bertschinger et al. [14] and the empirical methods provided in the IDTxL toolbox by Wollstadt et al. [15], as described in the methods.

Example activity from neuron pairs with high redundant (neurons R1 and R2) and synergistic (neurons S1 and S2) information components is shown in Fig 1(d) and (h), respectively. Both of these examples are taken from one session for a single mouse (mouse 4) in the Driscoll et al. [1] dataset. In these plots, the calcium signals ΔF/F of the two neurons are plotted against each other and colored according to the position of the mouse in the maze. The activity in each of these pairs is strongly correlated but in very different ways: in the high-redundancy pair, when one neuron is strongly activated, the other tends to be as well, whereas in the high-synergy pair, the two neurons are rarely activated together despite both being tuned to a position in the start of the maze. This correlation structure is largely representative of pairs with high redundancy and synergy in both datasets.

Fig 1(e) and (i) present a schematic view of how redundancy and synergy impact the pairs’ information about the position in the maze. In each plot, the ellipses represent contours of bivariate Gaussian distributions fit to the data in Fig 1(d) and (h). In the upper plots, the covariance matrices of the Gaussians are taken directly from the data, retaining their position-conditioned correlations. In the lower plots, the diagonal components of the covariance matrices are retained and the off-diagonal components are set to 0 to illustrate the activity of a pair of neurons with the same tuning and variance but no position-conditioned correlation. The purple and orange ellipses correspond to the activity in position bins 1 and 4 with the data binned into 7 position bins; only two bins are shown for the sake of visualisation. If these two ellipses overlap more, it is harder to determine the position of the animal given the activity of the two neurons. The effect of setting the covariance to 0 has opposite effects in the redundant and synergistic cases: in the former, there is less overlap than for the nonzero covariance, and in the latter there is more. This provides a rough picture of how neural correlations contribute to the information content of pairwise activity.

Fig 1(f) and (j) show the cumulative distributions of the redundant and synergistic components of the pairwise information over all neuron pairs for all mice in each dataset, with one distribution shown per behaviour or stimulus. These distributions have a very long tail, with the vast majority of pairs having very low redundant and synergistic information components.

### Redundancy index correlates with tuning stability

We asked whether a neuron’s redundant and synergistic interactions with the population are correlated with the stability of the neuron’s tuning to an external variable. In the two datasets we studied [1, 4], we found that the overall redundant coupling of a neuron to the population generally correlates with tuning curve stability.

We quantified a neuron’s degree of redundant and synergistic coupling with the population using the redundancy and synergy indices. For each neuron *x*_*i*_, we calculated and decomposed the joint mutual information between the external variable *u* and each pair containing that neuron {*x*_*i*_, *x*_*j*_}; *j* ∈ [1, *N*], *j*≠ *i*. This gives the four terms in Eq 2 for each pair of neurons containing *x*_*i*_. To obtain the redundancy and synergy indices Red and Syn, we averaged the redundant and synergistic contributions to the joint mutual information over all paired neurons *x*_*j*_, as

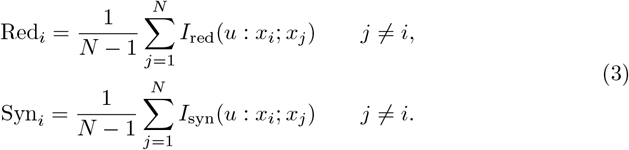

Fig 1(g) and (k) shows the cumulative distributions of the redundancy and synergy indices over all neurons in each dataset, for each behaviour and stimulus type. These distributions have long tails, similar to the redundant and synergistic information (Fig 1(f) and (j)), but with a less drastic drop in probability.

We then evaluated to what extent these indices were correlated with a neuron’s tuning stability. As a measure of tuning stability, we used the correlation-based representational drift index (RDI) defined by Marks et al. [4]:

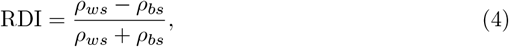

where *ρ*_*ws*_ and *ρ*_*bs*_ are the within- and between-session cross-correlation, respectively (see methods for details).

Neurons with high redundancy or synergy indices tend to be strongly tuned to the target variable. This is because the redundant and synergistic components of the pairwise mutual information are components of how two neurons together encode the stimulus, and when each neuron is taken individually, this information is still present in some form. To ensure any correlation we observed between these PID measures and the tuning stability were not simply due to this strong tuning, we included a measure of tuning strength—the single-neuron mutual information *I*(*u* : *x*) (Eq 1)—as a regressor. We constructed a linear model of stability using a multivariate regression with three regressors: mutual information (MI), redundancy index (Red), and synergy index (Syn). As shown in Fig 2(a), these regressors are all correlated with each other, so we used elastic net for regularisation.

**Fig 2.**
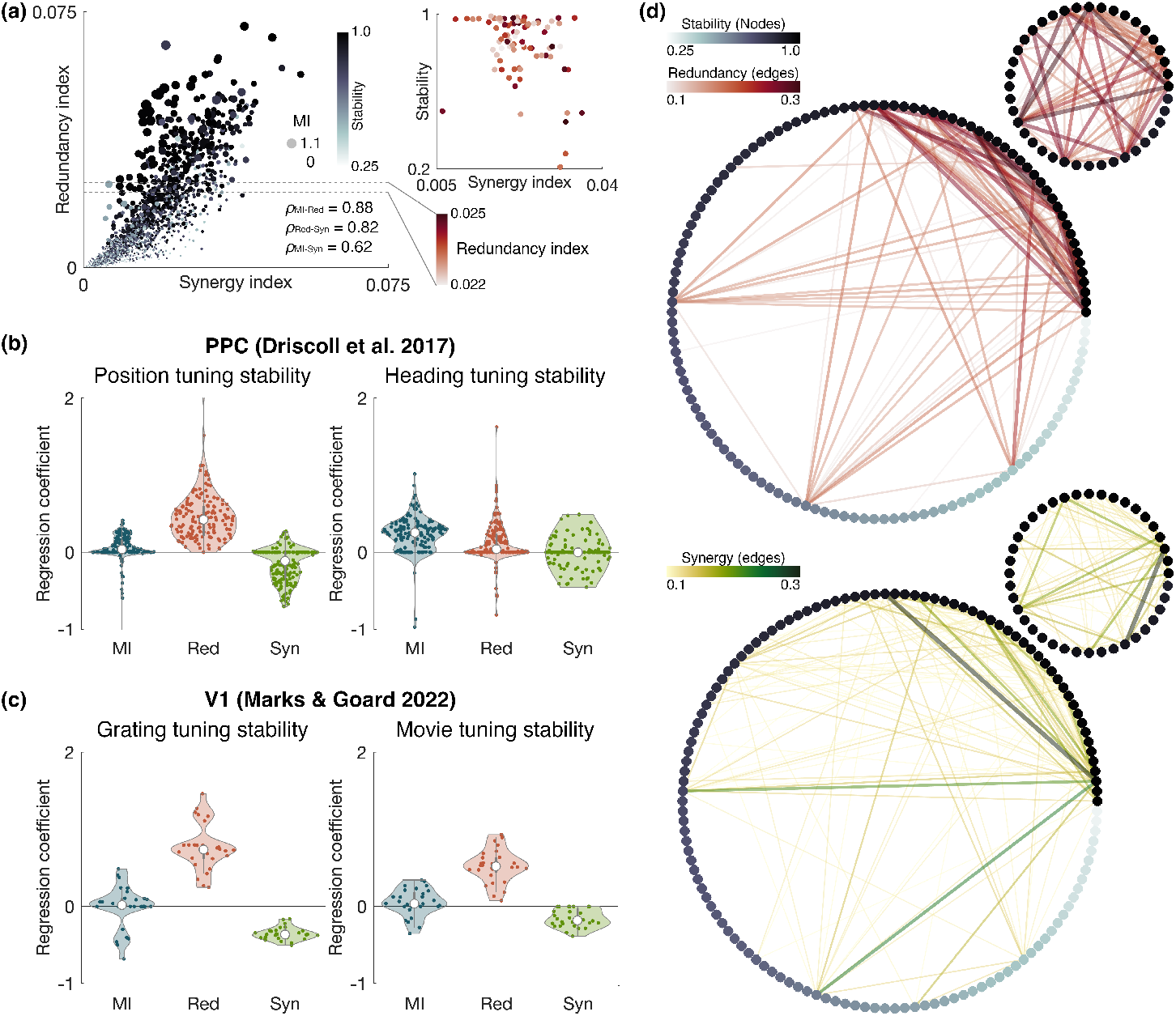
Redundancy index correlates with tuning stability. **(a)** Constructing a multivariate linear model for stability with three regressors: mutual information (MI), redundancy index (Red), and synergy index (Syn). The three regressors are all strongly correlated with each other. Each data point represents a neuron frome one mouse on a given day (Driscoll et al. [1], 130 neurons, 17 days). The redundancy index is plotted against the synergy index, with the size and colour showing the mutual information and stability, respectively. The session-averaged correlation coefficient *ρ* between each pair of regressors is given in the bottom right. **(b)** Regularised regression coefficients for the three predictors with position and heading as the target external variable [1]. Each data point represents a given session and trial type (left or right turns) for a given mouse. *n* = 617 PPC neurons across 4 mice. **(c)** Same as **(b)** but with gratings and movies as the external variable [4]. *n* = 1053 V1 neurons across 4 mice. **(d)** Graph plots showing pairwise synergy and redundancy across the network for one mouse on a single session (Driscoll et al. [1], 130 neurons, day 5, right turns). Each node represents a neuron, coloured and ordered according to position tuning stability, and each edge represents the pairwise redundant or synergistic component of the information, thresholded for visibility. Inset graphs show the top 40 most stable neurons in the population.

Fig 2(b) and (c) shows the regression coefficients for the three regressors for PPC tuning to position in the maze and heading of the animal and for V1 tuning to the drifting gratings and naturalistic movies, respectively. In most cases, the redundancy index was the most strongly weighted predictor of tuning stability. This indicates that the degree of redundant coupling to the population is a stronger indicator than the tuning strength of whether a neuron’s tuning will remain stable. The coefficient of the mutual information tended to be close to zero, indicating tuning strength provides little predictive power beyond that provided by the redundancy index.

Interestingly, the coefficient for the synergy index tended to be negative. This means that among neurons with a given redundancy index, a neuron with a higher synergy index would tend to exhibit lower tuning stability. A demonstration of this is shown in the inset in Fig 2(a). Here, the tuning stability is plotted against the synergy index for neurons within a narrow range of redundancy indices (0.022–0.025). Each data point in this plot corresponds to a neuron with a redundancy index in the target range on a given day and is colored according to redundancy index. As shown here, although the stability generally exhibits a positive trend with the synergy index, when the redundancy index is fixed, an underlying negative trend emerges.

One choice of external variable *u* did not follow these trends: tuning to the heading of the animal in the PPC. In this case, the mutual information was the most strongly weighted predictor of tuning stability, and the redundancy and synergy indices tended to be assigned coefficients near 0. This is most likely due to there being a sparser representation of heading among the recorded neurons. A pair of neurons must both be responsive to similar values of *u* to have any appreciable redundant or synergistic information components. Therefore, with a sparse representation, there is less chance for overlap in neurons’ tuning to *u*, making redundancy and synergy generally low across the population and therefore less predictive of stability. However, this result suggests that the relationship between stability and redundancy index is not a statistical artifact.

Network plots illustrating the pairwise redundancy and synergy on a single session for an example mouse from the Driscoll dataset are shown in Fig 2(d). In these plots, each node represents a single neuron, shaded according to its tuning stability, and the colour and thickness of each edge represents the redundant (top) or synergistic (bottom) component of the pairwise information. The small inset networks consist of the 40 most stable neurons in the population. As shown in the redundancy network, the most stable neurons are densely interconnected by high-redundancy edges, illustrating the relationship between redundancy and stability.

There are three neurons that appear to be exceptions to this: the three neurons to the left and near the bottom of the graph, with strong redundant connections to the most stable subset in the upper right quadrant. These neurons also have strong synergistic connections within the population, as shown in the bottom synergy graph. This further illustrates the interplay among redundancy, synergy, and stability indicated by the negative loading of the synergy index: neurons strongly coupled to the population by both redundant and synergistic pairings are less stable than those with predominantly redundant coupling.

## Discussion

The effects of representational drift are not seen uniformly across neurons in a population, and what contributes to this heterogeneity remains an open question. Although the results of Sweeney and Clopath [12] suggest neurons with more plastic representations couple more strongly to the population, our analysis is more aligned with other studies [9, 16] showing that neurons participating in functional ensembles tend to show greater tuning stability.

Here, we focused specifically on the effect of stimulus-conditioned correlations on information encoding. Our results demonstrate that the most stable cells in a population tend to participate in redundantly connected cliques that in turn synergistically connect to the rest of the population. Additionally, among these highly redundant subpopulations, those neurons with greater synergistic coupling to the population tend to be less stable (inset of Fig 2(a)). A closer look at select pairs of neurons suggests that redundant coupling may arise from either structural connections or tuning relationships among other unobserved variables. In either case, these relationships are indicative of a more complex population tuning to the task space not accounted for when considering a single neuron’s tuning to a single external variable [17].

On the basis of these observations, we conjecture that redundancy reveals how drift is constrained across the population in a way that limits interference with relationships among task variables that are more generally relevant across many tasks. Our observations here are somewhat limited by our discretisation of the task space and the confounding effect of the fixed relationships among the task variables. Further analysis using a different experimental paradigm and computational models would be beneficial to provide greater support for our findings. Despite these limitations, this work provides compelling evidence that stable subsets of the population encode for generally salient task features, evident in the stimulus-conditioned relationships among the neuronal responses. Furthermore, regardless of the mechanistic interpretation of the redundancy–stability relationship, our results also present a step toward a practical approach to predicting which neurons will maintain their relationship with the external variable, which is useful in technology reliant on long-term neural decoding such as brain–machine interfaces [13].

## Methods

### Experimental data

The two-photon calcium imaging and behavioural data used in this study are from Driscoll et al. [1] and Marks et al. [4], and complete details of the experiments and data can be found there.

#### Posterior parietal cortex

In Driscoll et al. [1], mice were trained over a 4–8 week period to perform a two-alternative forced-choice task in a virtual environment: the mice were given a visual cue at the start of a T-maze and had to turn left or right when they reached the T-intersection. Mice received a reward for correct associations of visual cues with left or right turns. The data are available from the Dryad repository (https://doi.org/10.5061/dryad.gqnk98sjq).

The MATLAB-based software ViRMEn (Virtual Reality Mouse Engine) [18] was used to construct the virtual environment and collect behavioural data. Driscoll et al. [1] collected calcium fluorescence imaging data at a sampling rate of 5.3 Hz and identified and segmented fluorescence sources. The processed fluorescence data used here consist of normalised fluorescence traces (Δ*F/F*).

In the present analysis, we excluded data from the inter-trial intervals and incorrect trials, and we analysed left- and right-turn trials separately, as many neurons display trial-type-specific activity [1]. Thus, the neuron response signal *x*(*t*) represents Δ*F/F* at time sample *t*, where *t* includes all time samples of all correct trials of a single type (i.e., left or right turn) on a given session. We considered two behavioural signals: position along the T-maze and heading angle.

Neurons had to be present (confidence index of 1 or 2) in at least 80% of the considered sessions (see Section) to be included in the analysis. We included four populations of neurons from three mice: mouse 3, 194 cells, 10 sessions (sessions 1, 2, 4, 6–12); mouse 3, 177 cells, 10 sessions (sessions 13–22); mouse 4, 130 cells, 17 sessions (sessions 1–17); mouse 5, 130 cells, 7 sessions (sessions 7–13).

#### Primary visual cortex

In Marks et al. [4], mice passively viewed two different types of visual stimuli: oriented gratings and a naturalistic movie. The gratings consisted of 12 different orientations evenly spaced from 0^°^ to 330^°^ in 30^°^ increments, presented in 8 trials. A single trial contained each orientation presented for 2 s followed by a 4-s inter-stimulus interval of a grey screen, with all orientations presented in ascending order of orientation angle. The naturalistic movie was a continuous 30-s clip from the film *Touch of Evil*, presented in 30 trials with a 5-s inter-trial interval between each presentation (grey screen). Mice viewed these stimuli once per week for 5–7 weeks; from the full dataset, we selected mice who had sessions for 7 weeks (mice 2, 10, 11, and 12 from the main experiment, mice 1–4 from the inhibitory experiment).

Marks et al. [4] collected calcium fluorescence imaging data at a sampling rate of 10 Hz and identified and segemented fluorescence sources. As with the data from Driscoll et al. [1], we performed our analysis on the normalized fluorescence traces (Δ*F/F*).

We excluded data from the inter-trial and inter-stimulus intervals, and we analysed the grating and movie trials separately. We constructed stimulus variables as the orientation angle of the gratings and the time elapsed in the movie.

As above, neurons had to be present (confidence index of 3) in at least 80% of sessions to be included in the analysis. We included four populations of excitatory neurons from four mice in the main experiment: mouse 2, 355 neurons; mouse 10, 252 neurons; mouse 11, 227 neurons; mouse 12, 219 neurons. We also analysed four populations of inhibitory neurons from four mice in the inhibitory experiment: mouse 1, 66 neurons; mouse 2, 76 neurons; mouse 3, 70 neurons; mouse 4, 48 neurons.

### Data discretisation

The nonparametric, information-theoretic estimators (mutual information, Δ*I*) used in this work require histogram-based estimators of various joint densities. To make this problem tractable, we discretised the fluorescence and behavioural signals prior to analysing the data (see [19] for details on data binning methods). Limitations of the data indicated a coarse bin size, which we verified was sufficient to support our conclusions.

We binned the fluorescence signals 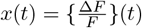 from each neuron into three unform-width bins. To reduce the influence of outliers, we first determined the 5^th^–95^th^ percentile range {*a, b*} = {percentile[*x*, 5], percentile[*x*, 95]}. We then defined the bin edges as 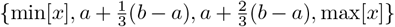. That is, we binned data within the 5^th^–95^th^ percentile range into three equally spaced bins, and assigned data below the 5^th^ or above the 95^th^ percentiles (saturation bounds) into the lower and upper bin, respectively.

The grating orientation stimulus variable from the Marks et al. [4] dataset is already a set of 12 discrete angles (0^°^ to 330^°^ in intervals of 30^°^) and thus requires no further binning. We binned the movie stimulus into 12 temporal bins of 2.5 s each.

We binned the behavioural signals from the Driscoll et al. [1] dataset (position and heading) into *N* = 5, 10, 20 state bins using two methods: uniform-width binning for quantifying tuning stability and uniform-activity binning for information theoretic metrics. Similar results were obtained for 10 and 20 bins.

We binned the heading into uniform-width bins in the same manner as with the fluorescence: with saturation bounds at the upper and lower fifth percentiles to account for extreme values. Because the position signal spans the T-maze track and is not influenced by extreme values in the same way as the other behaviours, we used the full range of position values to define the limits (*a, b*) for uniform-width binning. We used these uniform-width binned signals to calculate the change in tuning curves between a pair of days; this choice was made to ensure the bin edges were consistent across days.

In the Driscoll et al. [1] data set, neural population tuning is not distributed uniformly over all behavioural states when states are defined using the uniform-width approach. For example, more neurons respond near the beginning and end of the maze, and so the first and last position bins would be accompanied by higher population activity. For this reason, previous studies [19] have suggested partitioning covariates into bins containing (approximately) equal amounts of spiking activity. This results in higher-resolution binning for regions in behavior space that drive population activity more strongly.

In these data, we do not have direct access to spike counts, so we use the population-average Δ*F/F* as a proxy for this uniform-activity binning. For each behavioural variable *u*(*t*), we adaptively selected bin edges bins[*u*(*t*)] such that the integral of population-fluorescence activity was the same for each bin. That is, we (i) calculated the average total population activity for a given behaviour state *U* (over all correct left- or right-turn trials within a session), and (ii) adjusted the bin edges so that the sum of this average activity in each bin equaled the same constant *C*. Formally, this can be written as:

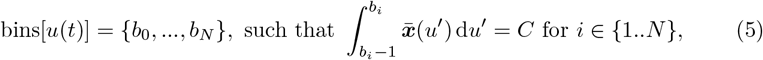

where 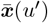 is the average population Δ*F/F* associated with the specific value *u*^*′*^ of the behavioral covariate. This is given by

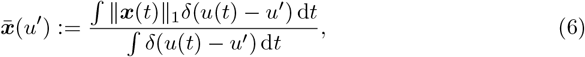

where ∥ ***x***(*t*) ∥_1_ is the sum of the population fluorescence activity for time-point *t*, and *δ* are Dirac deltas indicating times when the behavioral covariate *u*(*t*) has the specified value *u*^*′*^. We used the resulting uniform-activity binned signals to calculate the mutual information.

### Mutual information

We used the mutual information to quantify the strength of a neuron’s tuning to a target behavioural variable. The mutual information *I*(*u* : *x*), in bits, between a neuron response *x*(*t*) and a behaviour *u*(*t*) is given by

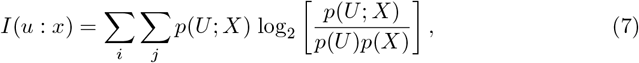

where *U* and *X* are the state values of the discretised behaviour *u* and response *x* vectors, respectively, and *p*(·) denotes the probability.

We used a shuffle test to determine whether each neuron was significantly tuned to the target behaviour. For this test, we shuffled the time samples of the concatenated within-trial neuron responses 1000 times for each neuron and recalculated the mutual information between the behaviour and the shuffled signals. If the mutual information of the unshuffled signal is greater than 95% of the shuffled cases, the neuron is considered tuned to the behaviour.

### Quantifying synergy and redundancy

To quantify how stimulus-conditioned correlations between pairs of neurons influence the information the pair encodes about an external variable *u*, we decomposed the pairwise mutual information into its constituent components. The mutual information *I*(*u* : *x*_1_, *x*_2_) between an external variable *u* and a pair of neurons {*x*_1_, *x*_2_} can be decomposed into four terms, as

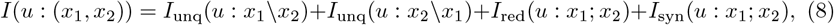

where *I*_unq_ is the information provided uniquely by one of the neurons (unique information), *I*_red_ is the information both neurons share about *u* (redundant information), and *I*_syn_ is the information they provide together (synergistic information).

Various methods of estimating the terms in Eq 8 have been proposed. In this study, we used the formulation by Bertschinger et al. [14] and the empirical methods to compute them provided in the IDTxL toolbox by Wollstadt et al. [15]. Following Bertschinger et al. [14], the four components of the information can be estimated from data as

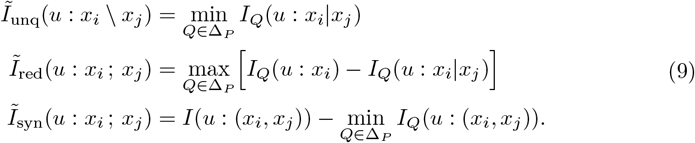

In the notation used in Eqs. 8 and 9, a colon between variables indicates the mutual information is computed between them, a backslash indicates information present in the former variable but not the latter (unique information), and a semicolon indicates information related to the interaction of the two variables (redundant or synergistic information).

Eq 9 involves maximising or minimising information terms *I*_*Q*_ across the probability distributions *Q* in the set of distributions Δ_*P*_. Letting Δ be the set of all joint distributions of the three variables of interest {*u, x*_*i*_, *x*_*j*_}, Δ_*P*_ is the subset of Δ in which the marginal distributions on the pairs (*u, x*_*i*_) and (*u, x*_*j*_) are the same as in the empirical distribution *P* :

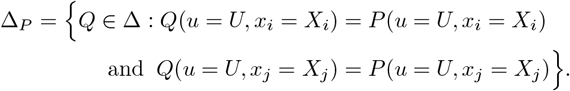

As in the definition of the mutual information, capital letters *U* and *X* represent state values of the discretised signals *u* and *x*. We performed the optimisations in Eq 9 using the method by Makkeh et al. [20], as implemented in IDTxL [15].

To characterise the tendency of a neuron to interact redundantly and synergistically with the population, we defined redundancy and synergy indices for each neuron. The redundancy and synergy indices Red and Syn are obtained by averaging the redundant and synergistic contributions to the joint mutual information over all paired neurons *x*_*j*_, as

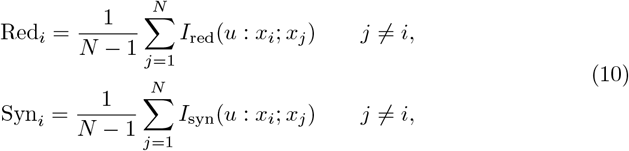

where *N* is the number of neurons in the population.

### Quantifying tuning stability

We quantified the tuning stability of each neuron by comparing its tuning curves between pairs of sessions. To evaluate the robustness of the observed correlations to methodological choices, we used three tuning curve comparison metrics—(1) the cross-correlation [4], (2) the cosine similarity [21], and (3) the mean of absolute differences. The main results are reported using the cross-correlation–based method, for the sake of consistency with the original results from Marks et al. [4]. However, all three stability methods yielded results consistent with those shown in Fig 2. Each of the three methods are described in more detail below.

First, we normalised the tuning curves as follows. The raw session-wise tuning curves are given as an *N* -dimensional vector, where *N* is the number of behavioural bins, and each component in the vector is the average Δ*F/F* across all samples in the given behavioural bin. These vectors are then z-scored to give the normalised tuning curve.

The cross-correlation metric of stability is given by

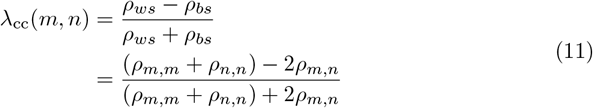

where 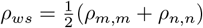 and *ρ*_*bs*_ = *ρ*_*m,n*_ are the Pearson correlations within and between sessions. To calculate the within-session correlation, the trials are randomly partitioned in half and the correlation is taken between the two halves. This partitioning is performed 1000 times and the final within-session correlation is the average over the 1000 partitions. The overall tuning stability for the neuron is then given by Λ_*cc*_ = ⟨*λ*(*m, n*) ⟩_*m,n*; *m ≠ n*_, the average over all pairs of sessions, as is also the case for the other two metrics.

The cosine similarity metric of stability [8] is given by

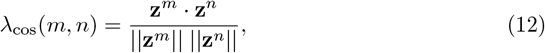

where **z**^*m*^ and **z**^*n*^ are the normalised tuning curves on sessions *m* and *n*, respectively. This metric is then normalised by the within-session cosine similarity during each session *m* and *n*. As with the cross-correlation metric, the within-session stability is averaged over 1000 random partitions. This approach considers the ‘alignment’ of the tuning curve vectors in *N* -dimensional space and can range from 0 (orthogonal) to 1 (perfectly aligned).

For the mean of absolute differences method, the tuning stability *λ*(*m, n*) for a pair of sessions *m* and *n* (*m*≠ *n*) is given by

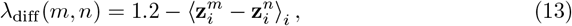

where 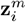 and 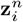 are the *i*th components of the normalised tuning curves on sessions *m* and *n*, respectively. The subtraction from 1.2 is included to yield stability values that are increasingly positive for increasingly similar tuning curve pairs; the selection of the constant value was chosen to yield stability values generally falling in the range of 0 to 1.

### Statistical relevance of predictors of drift

A primary goal of this paper was to determine whether redundancy and synergy are predictive of a neuron’s stability. We used regularised regression, specifically elastic net, to evaluate which predictor variables contribute to describing variance in stability and rank their predictive relevance. This regularisation method is useful when handling linear models with strongly multicollinear predictors, as it penalises high coefficients to avoid overfitting. The resulting coefficients can thus be used to rank the relevance of the corresponding predictors in the model.

We used three predictor variables in our regression model: the redundancy index, the synergy index, and the single-neuron mutual information. The three predictor variables and the tuning stability were all z-scored before the regression. If any predictor did not have a significant correlation with the stability in a single-variable model (i.e., *p >* 0.05), its regression coefficient was set to 0. Elastic net combines the L1 penalty of lasso regression with the L2 penalty of ridge regression, and we used a mixing parameter of *α* = 0.5 to define the loss function. We selected the *λ* parameter corresponding to the model with the lowest mean squared error to obtain the final coefficient values.

## Acknowledgments

This project has received funding from the Research Council of Norway (NFR) IKTPLUSS grant (SOCRATES, grant no. 270961, awarded to KH and SN), the NFR FRIPRO grant (DeepCA, grant no. 286558; awarded to SN), the HORIZON EUROPE European Research Council (ERC) starting grant (FLEXNEURO, grant no. 716643, awarded to TO), the Human Frontier Science Program (HFSP) grant (grant no. RGY0069, awarded to TO), the Leverhulme Trust fellowship (awarded to MER), the Isaac Newton Trust fellowship (ECF-2020-352, awarded to MER), the Alexander von Humboldt postdoctoral fellowship (awarded to KH), and the Theoretical Sciences Visiting Program (TSVP) at the Okinawa Institute of Science and Technology (OIST) (awarded to TO). The funders had no role in study design, data collection and analysis, decision to publish, or preparation of the manuscript.

We would like to thank Joseph Lizier and Michael Goard for insightful discussions about this work.

